# Construct a molecular associations network to systematically understand intermolecular associations in *Human* cells

**DOI:** 10.1101/693051

**Authors:** Hai-Cheng Yi, Zhu-Hong You, Zhen-Hao Guo

**Affiliations:** Xinjiang Technical Institute of Physics and Chemistry, Chinese Academy of Sciences, Urumqi 830011, China; University of Chinese Academy of Sciences, Beijing 100049, China

## Abstract

A key aim of post-genomic biomedical research is to systematically integrate and model all molecules and their interactions in living cells. Existing research usually only focusing on the associations between individual or very limited type of molecules. But the interactions between molecules shouldn’t be isolated but interconnected and influenced. In this study, we revealed, constructed and analyzed a large-scale molecular association network of multiple biomolecules in *human* cells by modeling all associations among lncRNA, miRNA, protein, circRNA, microbe, drug, and disease, in which various associations are interconnected and any type of associations can be predicted. More specifically, we defined the molecular associations network and constructed a molecular associations dataset containing 105546 associations. Then, each node is represented by its attribute feature and network embedding learned by Structural Deep Network Embedding. Moreover, Random Forest is trained to predict any kind of associations. And we compared the features and classifiers under five-fold cross-validation. Our method achieves a remarkable performance on entire molecular associations network with an AUC of 0.9552 and an AUPR of 0.9338. To further evaluate the performance of our method, a case study for predicting lncRNA-protein interactions was executed. The experimental results demonstrate that the systematic insight for understanding the synergistic interactions between various molecules and complex diseases. It is anticipated that this work can bring beneficial inspiration and advance related systems biology and biomedical research.

**Author Summary:** The interactions between the various biomolecules in the cells should not be isolated, but interconnected and influenced. There have been many valuable studies on the interactions between two individual molecules. Based on a systematic and holistic perspective, we revealed and constructed a large-scale molecular associations network by combining various associations in human living cells, including miRNA-lncRNA association, miRNA-disease association, miRNA-protein interaction, lncRNA-disease association, protein-protein interaction, protein-disease association, drug-protein interaction, drug-disease interaction, and lncRNA-protein interaction. To model and analyze this molecular associations network, we employed the network representation learning model to learn how to represent the node. Each node in the network can be represented by network embedding and its own attribute information. Any node can be classified. And any type of the associations in this network can be predicted, which can be considered as link prediction task. Our work provides a new systematic view and conceptual framework to understand complex diseases and life activities. It is anticipated that our study can advance related biological macromolecules, systems biology and biomedical research, and bring some meaningful inspiration.

## Introduction

In the past century, reductionism alternately guide the development of biology, has provided a wealth of valuable knowledge and data resources on individual cellular components and their functions. But the study of isolated cellular components does not help to answer the core questions about complex life activities, such as the nature of cells and the essence of life. A key goal of post-genomic biomedical research is to systematically integrate and model all molecules and their interactions in living cells. Understanding these molecules and their associations, whether isolated or surrounded by other cells, is essential to understanding how they determine the functions of this extremely complex machine of cells. As a master of reductionism theory, molecular biology and genomics have thoroughly pursued the fine structure and underlying physical basis of cellular system components, and accumulated excessive experimental data resources. Comprehensive and accurate modeling and understanding of these data requires a systematic and holistic perspective and approach. As a new holism, system biology inherits existing research paradigms such as molecular biology and genomics, providing a new understanding and framework for our study of life activities and complex diseases.

Existing biomolecular interaction or association studies often focusing only on the links between individual molecules, including mRNA-protein interactions [1, 2], lncRNA-protein interactions [3], protein-protein interactions [4], miRNA-protein interactions [5], miRNA-lncRNA interactions [6, 7], considering exogenous chemical compound or complex disease, there is drug-protein interactions [8, 9], drug-disease interactions [10-12], miRNA-disease associations [13, 14], lncRNA-disease associations [15], protein-disease associations [10, 16]. Emerging research on circRNA shows there are also circRNA-miRNA associations [17], circRNA-protein interactions [18] and circRNA-disease associations [19]. From a systematic perspective, as one of the participating elements of human life activity system, the environment, or simply, the microbes, also has an association with disease [20] and affects the functions of drug [21]. In this study, we releveled and defined these interconnected associations between biomolecules as the Molecular Associations Network (MAN). The molecular associations in the *human* are shown in the Figure 1:

**Figure 1.**
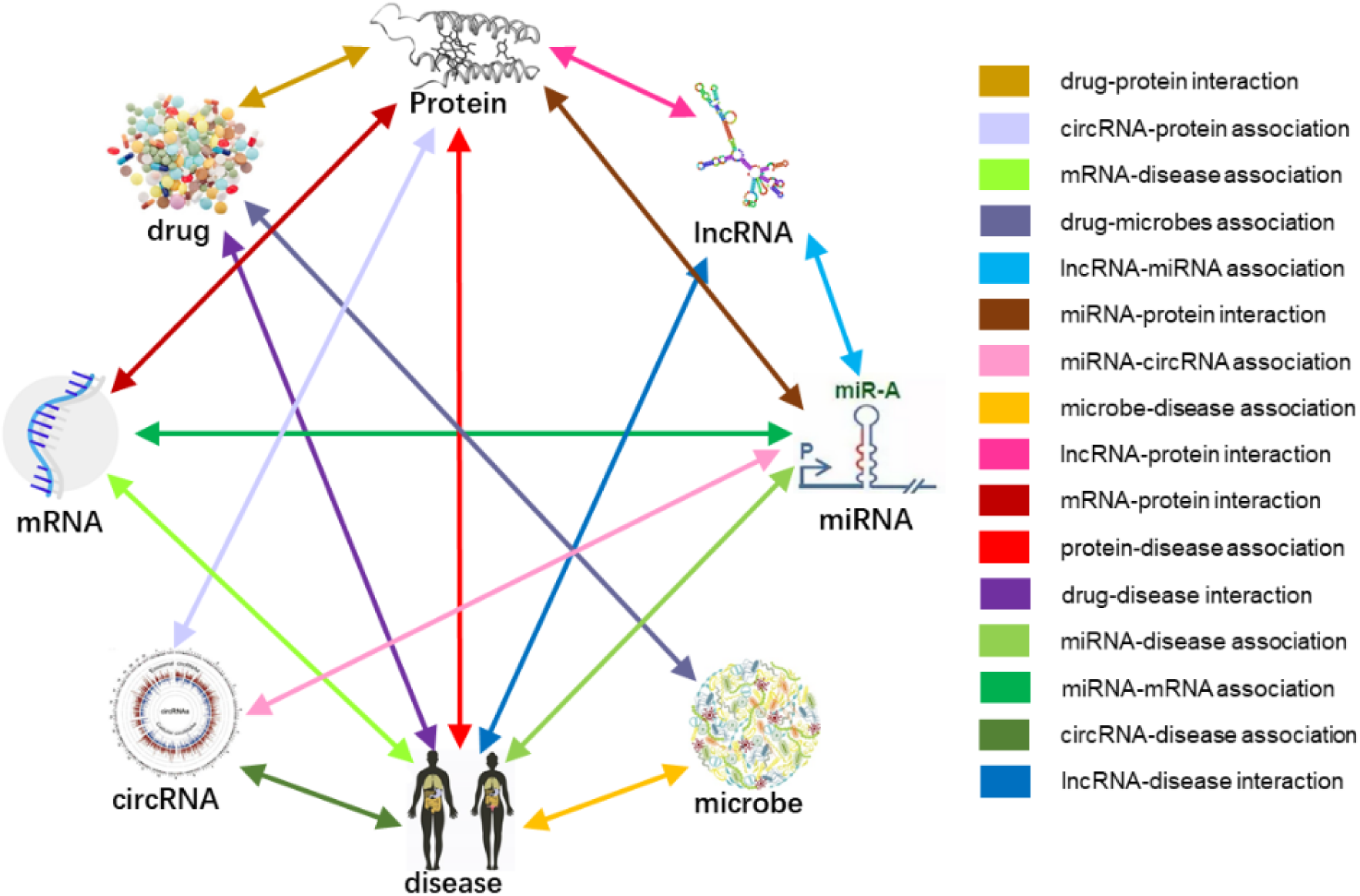
The molecular associations exist in *human*.

There are several studies have considered the association between more than one biomolecule. Davis *et al*. manually compiled chemical-gene interactions, chemical-disease relationships, and gene-disease relationship data from literature, which provides a wealth of information for integrating data to constructed a chemical-gene-disease network [22]. Liu *et al*. constructed a heterogeneous network to predict miRNA-disease associations by connecting miRNA-target gene, miRNA-lncRNA associations, and miRNA-disease associations to calculated miRNA similarity and disease functional similarity [23]. Similarly, Chen *et al*. used lncRNA as a mediator to discover the associations between miRNA and disease by constructing a heterogeneous network of miRNAs, lncRNAs and diseases [24]. Zhu *et al*. described a Drug-Gene-ADR heterogeneous information network and use it to predict drug-gene interactions [25]. Lin *et al*. studied multimodal networks linking diseases, genes and chemicals (drugs) by applying three diffusion algorithms and different information content [26]. Yang *et al*. integrated protein-protein interactions and cell lines’ functional contexts similarity to predict drug response [27]. But the types of biomolecules integrated in these studies are still very limited. Moreover, these studies are not based on the perspective of systematic intermolecular interactions.

Network is an important form to express the connection between objects and objects. A key problem in the analysis of networks is to study how to reasonably represent the characteristic of nodes in the network. In order to construct and analyze large-scale heterogeneous molecular associations network, we need appropriate network representation methods. Early matrix eigenvectors-based network representation learning algorithms strongly depend on the construction of relation matrices, for example, locally linear embedding [28], Laplace eigenmap [29] and directed graph embedding [30]. Such methods generally first define a linear or second-order loss function for the representation of the node. Then the optimization problem is transformed into the eigenvector calculation problem of a relation matrix. The main disadvantage of this type of method is the computational complexity: the eigenvector calculation of large-scale matrices is very time and space consuming. Neural network-based methods have made remarkable progress in the field of natural language and image processing. Similarly, such methods are also effective in the field of network representation [31]. For the first time, DeepWalk [32] introduced neural network model into the network representation learning field using Random Walk and word2vec, making full use of the information network structure. Another representative network representation learning algorithm is LINE [33]. It models all pairs of first-order similarity and second-order similarity nodes and minimizes the KL distance between the probability distribution and the empirical distribution, which can be applied to large-scale directed weighted graphs network representation learning. In addition, matrix factorization is also an important means of network representation learning [34, 35]. Structural Deep Network Embedding (SDNE) [36] is applied to learn representation of each node in MAN, namely MAN-SDNE.

In this study, we defined a large-scale molecular association network of multiple molecules associations in *human* cells and constructed a dataset of MAN. It provides a systematic perspective for understanding the synergistic interactions between various molecules and complex diseases. More specifically, we cleaned and collected the most widely available data sets of nine biomolecular interactions to construct the whole molecular associations dataset (MAD). These data sets are originally isolated associations among five different objects, including miRNA, lncRNA, protein, drug and disease, with redundancy and duplicate names between entries, we unify it into a unified naming system and remove redundancy. Meanwhile, we introduce the network representation learning algorithm SDNE into MAN to calculate feature representation of each node in the MAN. And then, we simplify the heterogenous network to be homogeneous. Moreover, we consider not only the network embedding features, but also the attribute characteristics of the nodes themselves, such as k-mer frequency of lncRNA, miRNA, protein sequence, Morgan fingerprint of drug compound structure and semantic similarity of disease phenotype. All known association pairs are positive samples, and the same number of unknown pairs that are randomly selected as negative samples to form a data set. Random Forest (RF) is used as the classifier for link prediction, and to avoid bias, the RF uses only default parameters and does not make any adjustments. Five-fold cross-validation was adopted to verify the performance of the model. Furthermore, we compared the performance of different features and different classifiers. And a case study using MAN-SDNE to predict lncRNA-protein interactions was carried out. The experimental results are in line with expectations, the MAN contained rich information and MAN-SDNE can predict whether any node pairs have interactions. The reveal of MAN offers a new systematic view of complex diseases and life activities. It is anticipated that this research could advance systems biology and biomedical research, or bring some useful inspiration.

## Materials and Methodology

### Construction of Molecular Associations Dataset

In the past few decades, the interactions between individual biomolecules have been well studied. Some existing research provides valuable data on individual molecular interactions or associations. In order to create a comprehensive dataset of MAN, we have collected the most extensive data sets for various molecular associations in *human* cells, including miRNA-lncRNA from lncRNASNP2 [37], miRNA-disease from HMDD [38], miRNA-protein from miRTarBase [39], lncRNA-disease from LncRNADisease [40] and lncRNASNP2 [37], lncRNA-protein from LncRNA2Target [41], protein-disease from DisGeNET [42], drug-protein from DrugBank [43], drug-disease from CTD [44], and protein-protein from STRING [45]. In these data sets, the same entity may have different naming conventions. To connect different interactions of the same entity, we unify them into a unified naming scheme. Further, we remove redundancy and duplication. Finally, we obtained a MAD of 105546 association records with five kinds of molecules objects, nine types of interactions. The details of node types and associations items in MAD are shown in Table 1 and Table 2 below.

**Table 1.** The amount of 5 types of nodes in the MAD dataset.

| Node type | Amount |
| --- | --- |
| **lncRNA** | 769 |
| **miRNA** | 1023 |
| **protein** | 1649 |
| **drug** | 1025 |
| **disease** | 2062 |
| **Total** | 6528 |

**Table 2.** Type of association, source and quantity in the MAD.

| Association type | Amount | Database |
| --- | --- | --- |
| miRNA-lncRNA | 8374 | lncRNASNP2 |
| miRNA-disease | 16427 | HMDD |
| miRNA-protein | 4944 | miRTarbase |
| lncRNA-disease | 1264 | LncRNADisease<br>lncRNASNP2 |
| Protein-protein | 19237 | STRING |
| Protein-disease | 25087 | DisGeNET |
| Drug-protein | 11107 | DrugBank |
| Drug-disease | 18416 | CTD |
| lncRNA-protein | 690 | lncRNA2Target |
| Total | 105546 | This study |

### Network embedding of the nodes

To obtain high effective node features for large-scale network, the network representation learning model SDNE is adopted to learn low-rank representations of each node and capture network structure. Unlike previous methods of using shallow neural networks, SDNE uses deep neural networks to model the nonlinearity between node representations. The entire model can be divided into two parts: one is model the first-order proximity supervised by the Laplace matrix. The other is to model the second-order proximity by an unsupervised deep autoencoder. The final SDNE algorithm uses the output of middle layer of the deep autoencoder as the network representation of the node. The framework of SDNE is shown in Figure 2.

**Figure 2.**
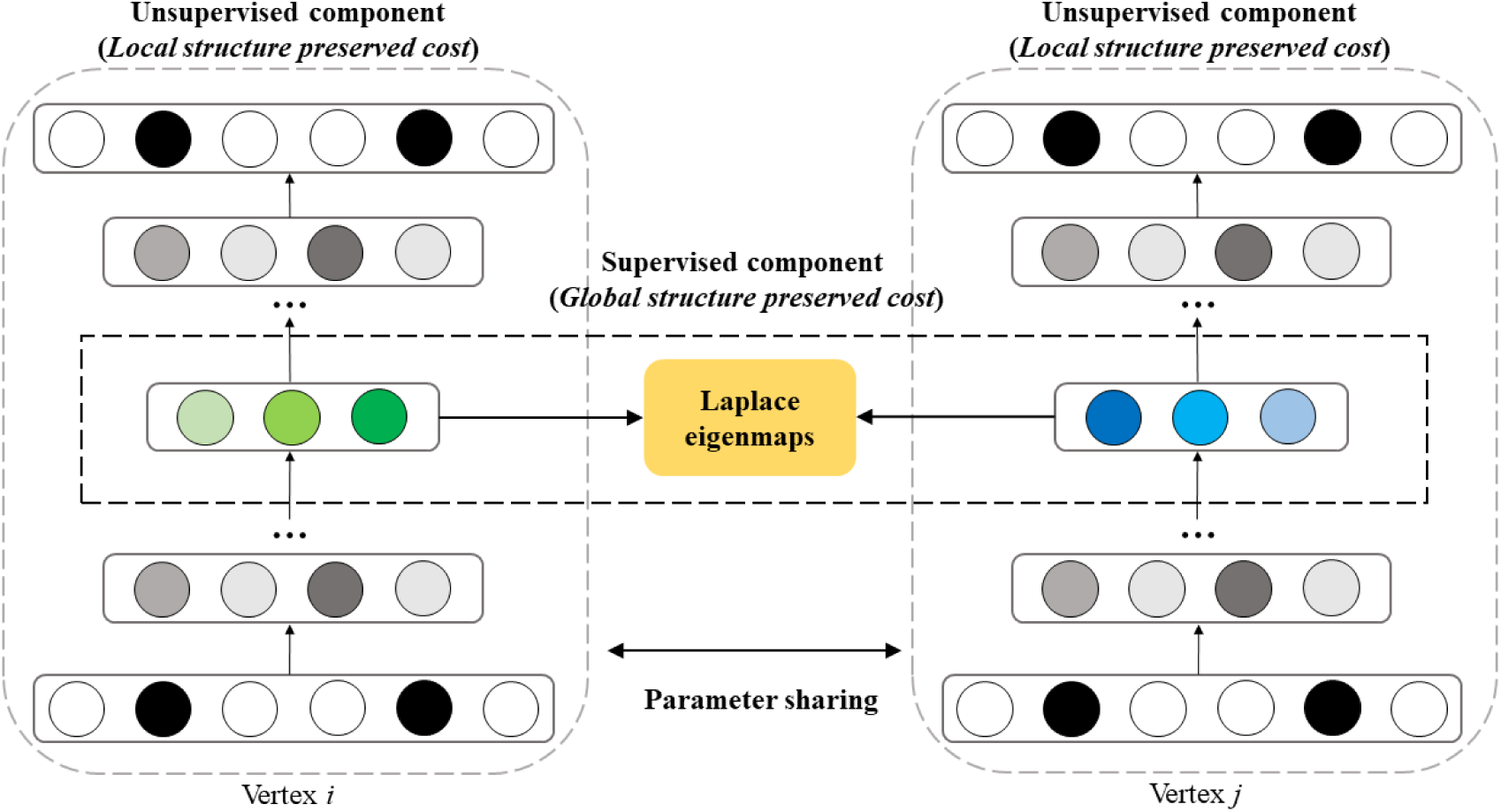
The framework of the semi-supervised deep model SDNE.

The details of second-order proximity that unsupervised components use to preserve global network structures was first described. Suppose a network *G =* (*V, E*), its adjacency matrix *M*, which contains *n* instances *m*_*1*_, *m*_*2*_, …, *m*_*n*_. for each instance m_i_, it can be defined as:

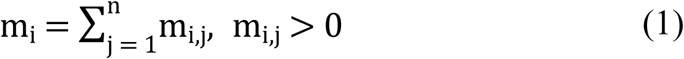

Only if there is a link between *v*_*i*_ and *v*_*j*_. while *m*_*i*_ means the neighborhood structure of the node *v*_*i*_ and *M* contains the neighborhood structure of each node. SDNE applied the deep autoencoder to obtain the second-order proximity.

The objective function can be defined as follows:

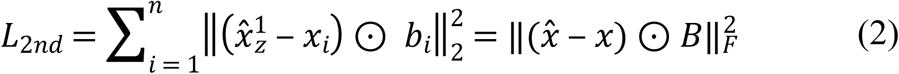

Where ⊙ indicates the Hadamard product. SDNE not only considers the global network structure, but also the local structure, which is represent by first-order proximity. The first-order proximity can be viewed as the supervised information to constrain the similarity of the potential representations of a pair of nodes. The loss function of the first-order proximity can be defined as:

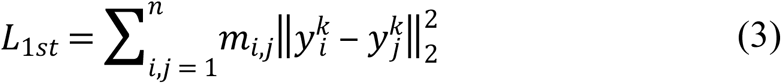

The final semi-supervised model SDNE combines the first-order and second-order proximity simultaneously. The objective function is shown as below:

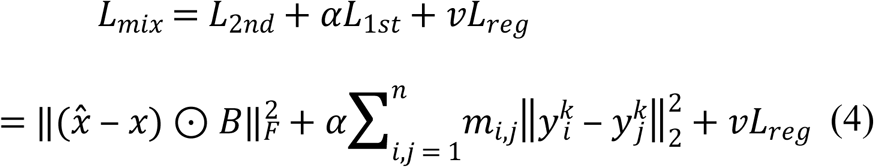

where *L*_*reg*_ is an *L*_2_-norm regularization term to avoid overfitting. The define of *L*_*reg*_ is:

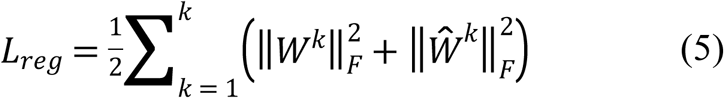

### Node attributes

Each node in the MAN can be defined not only the network embedding, but also the attribute of themselves. In this work, for node with sequence information, the k-mer frequency was applied to exploit their attribute feature. For drug, Morgan fingerprints that represent their chemical structure are used as attribute features. For disease, we use a Medical Subject Headings (MeSH) descriptor describing the phenotype of the disease to construct a directed acyclic graph (DAG) to calculate disease similarity, using this measure of similarity as the attribute of the disease.

For sequence of miRNA, lncRNA, we use the 3-mer frequency to encode its sequence, from AAAA to UUUU, there is 4^*k*^ possible combinations of nucleic acid residues (A, C, G, U). For a given sequence, slide from left to right four residues as a sliding window, one residue one step, we can obtain the composition information of a sequence, and then, we normalize the feature vector according to the sequence length.

For protein, the processing of protein sequences is slightly different. The 20 amino acids are first divided into four groups according to the polarity of the side chain, which is inspired by existing protein study [46], including (Ala, Val, Leu, Ile, Met, Phe, Trp, Pro), (Gly, Ser, Thr, Cys, Asn, Gln, Tyr), (Arg, Lys, His) and (Asp, Glu). Then we can use the same 3-mer frequency mentioned above to process the protein sequence.

For drug, the chemical structure is represented by Simplified Molecular Input Line Entry Specification (SMILES) [47], then we calculate corresponding Morgan fingerprints for each compound.

For disease, we use DAG to represent each disease based on MeSH descriptor. DAG(*D*) = (*D, N*_(*D*)_, *E*_(*D*)_), *N*_(*D*)_ is the set of points that contains all the diseases in the DAG(*D*). *E*_(*D*)_ is the set of edges that contains all relationships between nodes in the DAG(*D*). For the diseases that are included in MeSH, the semantic similarity that is calculated by means of DAG can be chose to represent the disease according to the previous literature. The semantic similarity between different diseases can be defined as follows. In DAG of disease *D*, the contribution of any ancestral disease *t* to disease *D* is as the formula:

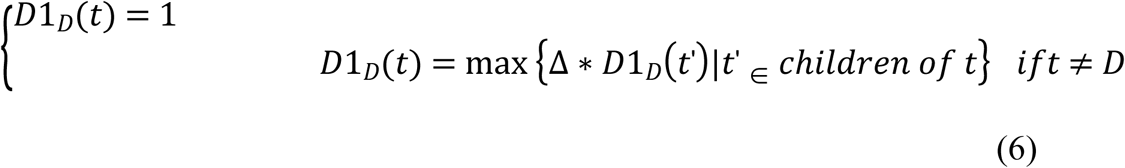

Δ is the semantic contribution factor. The contribution of disease *D* to itself is 1 and the contribution of other nodes to *D* will be attenuated due to Δ. Based on Equation (1), we can obtain the sum of the contributions of all diseases in DAG to *D*:

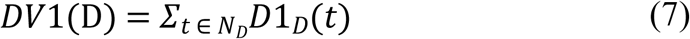

Like the Jaccard similarity coefficient, the semantic similarity between the diseases *i* and *j* can be calculated by the following formula:

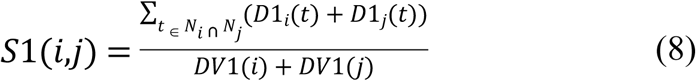

Considering that the dimensions of attribute feature vector of different kinds of nodes are not uniform, we trained a deep autoencoder to learn its hidden high-level low-rank representation and unify its dimensions.

### Deep autoencoder

In the framework of abovementioned SDNE, deep autoencoder (DAE) was used. For self-contained, in this section, we will briefly review the core ideas of DAE. It is an unsupervised deep learning model consisting of two parts: encoder and decoder. The encoder consists of several nonlinear functions that map the input data to the representation space. The decoder includes a plurality of non-linear functions that map representations in the representation space to the reconstruction space. For a given input *x*, DAE maps the input to the output *O*(*x*):

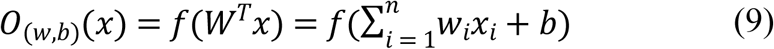

where the nonlinear activation function *f* can be defined as:

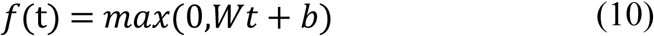

Suppose the output of *O*(*x*) is 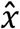, DAE aims to minimize the error between input and output. The loss function can be defined as follow:

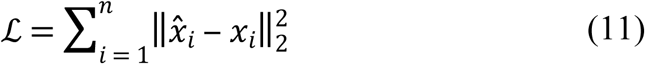

### Performance evaluation

In this study, the widely used evaluation measure was followed to evaluate our method, including accuracy (Acc.), sensitivity (Sen.), also means recell, specificity (Spec.), precision (Prec.) and Matthews Correlation Coefficient (MCC) defined as:

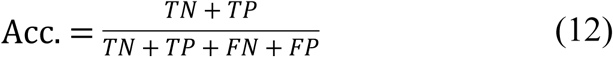

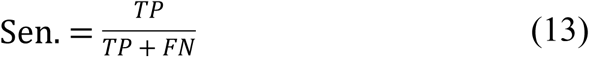

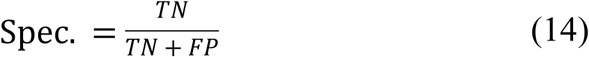

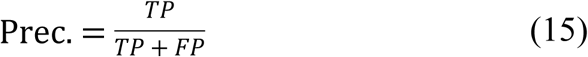

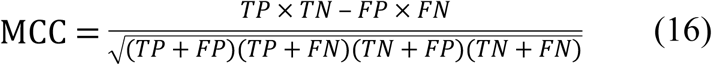

where *TN* represents the correctly predicted number of negative samples, *TP* stands for the correctly predicted number of positive samples, *FN* indicates the wrongly predicted number of negative samples and *FP* denotes the wrongly predicted number of positive samples. Certainly, the Receiver Operating Characteristic (ROC) curve, Precision-Recall curve and the area under the ROC curve (AUC), the area under the precision-recall curve (AUPR) are also adopted to evaluate the performance of MAN-SDNE.

## Results and Discussion

### Five-fold cross validation performance of MAN-SDNE

First, we will briefly introduce the scheme of five-fold cross validation. The entire data set is randomly divided into five equal parts, each taking four subsets as the training set and the remaining one subset as the test set, cycle five times in turn, take the average of five times as the final performance. For MAN, we remove 20% of the links each time as the training set, the removed links as the testing set, and ensure that the links removed in these five times have no overlap. In each fold cross validation, we only use the training set as input to the network representation learning model to learn the network embedding of nodes. The five-fold cross validation performance is shown in Table 3 and Figure 3 as follow:

**Table 3.**
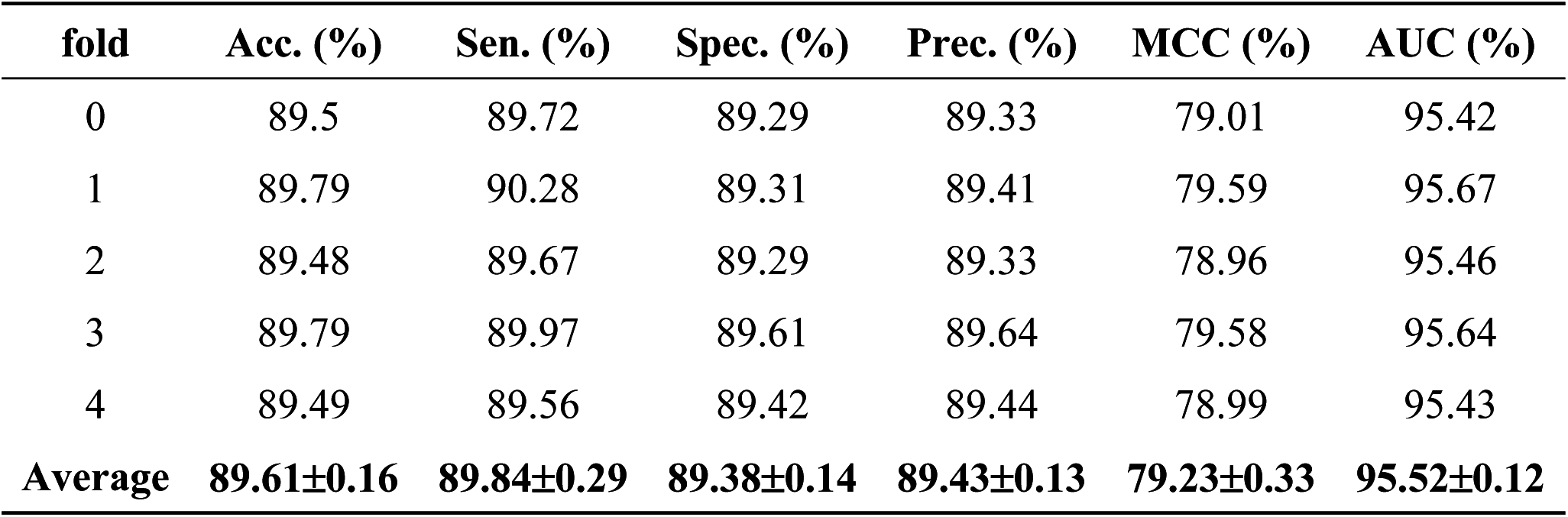
The five-fold cross-validation performance of MAN-SDNE on the entire MAN dataset.

| fold | Acc. (%) | Sen. (%) | Spec. (%) | Prec. (%) | MCC (%) | AUC (%) |
| --- | --- | --- | --- | --- | --- | --- |
| 0 | 89.5 | 89.72 | 89.29 | 89.33 | 79.01 | 95.42 |
| 1 | 89.79 | 90.28 | 89.31 | 89.41 | 79.59 | 95.67 |
| 2 | 89.48 | 89.67 | 89.29 | 89.33 | 78.96 | 95.46 |
| 3 | 89.79 | 89.97 | 89.61 | 89.64 | 79.58 | 95.64 |
| 4 | 89.49 | 89.56 | 89.42 | 89.44 | 78.99 | 95.43 |
| **Average** | **89.61±0.16** | **89.84±0.29** | **89.38±0.14** | **89.43±0.13** | **79.23±0.33** | **95.52±0.12** |

**Figure 3.**
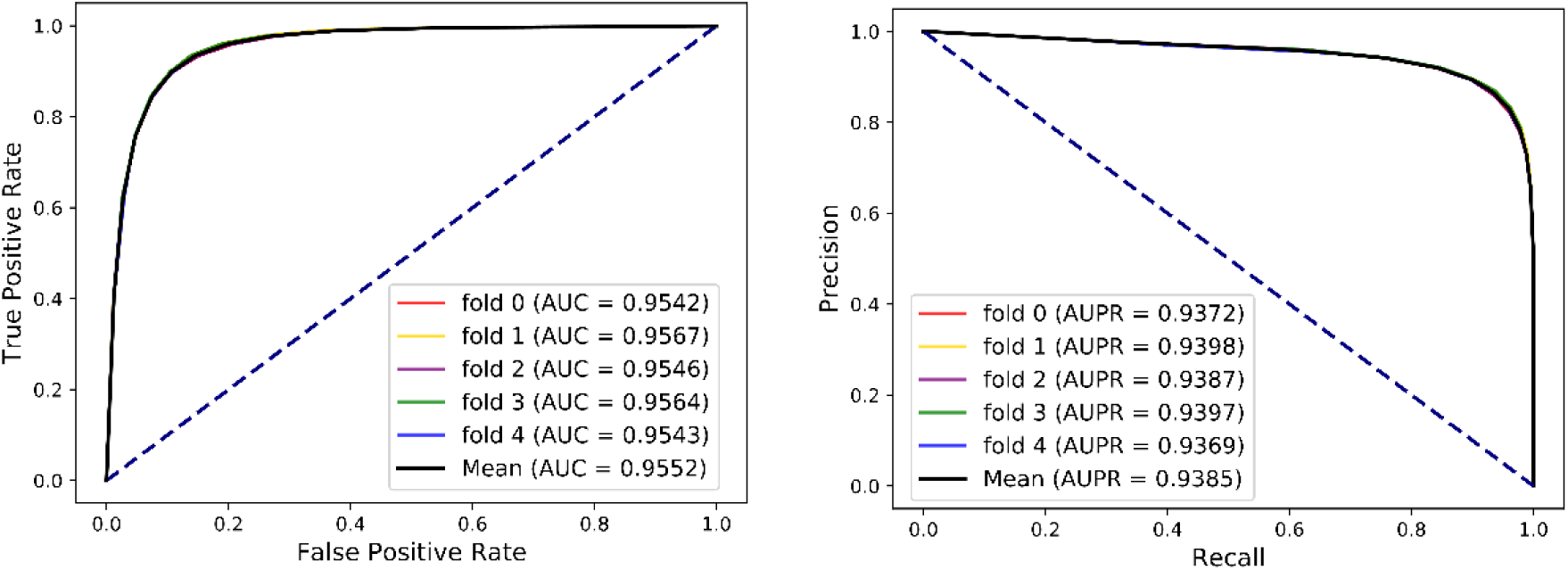
The five-fold cross validation ROC, Precision-Recall curve, AUC and AUPR of MAN-SDNE on the entire MAN dataset.

On entire MAN, for predicting any type of molecular associations, that is, for predicting any link or edge in the associations network, our method MAN-SDNE achieves an average accuracy of 89.61%, a sensitivity of 89.84%, a specificity of 89.38%, a precision of 89.43%, a MCC of 79.23%, a AUC of 95.52% and a AUPR of 0.9385. It should be noted that our classifier only uses the default parameters and does not perform any parameter optimization. To characterize the volatility of the model’s performance, we also calculated the standard deviation of the five-fold cross-validation. As can be seen from Table 3, the standard deviation of the above indicators is 0.16, 0.29, 0.14, 0.13, 0.33, and 0.12, which can reflect that our model MAN-SDNE is very stable and robust.

### Evaluate and compare the effects of network embedding and attribute features

To fully exploit the discriminative features of nodes, we considered both the network embedding and the attribute of nodes in the MAN. In this section, we will evaluate and compare the effects of individual network embedding and attribute features and their combined use. The details are shown in Table 4 and Figure 4 as follow.

**Table 4.** Comparison of different features.

| Feature | Acc. (%) | Sen. (%) | Spec. (%) | Prec. (%) | MCC (%) | AUC (%) |
| --- | --- | --- | --- | --- | --- | --- |
| Attribute | 87.92±0.2 | 90.51±0.33 | 85.33±0.35 | 86.06±0.27 | 75.95±0.4 | 93.8±0.11 |
| Embedding | 87.19±0.43 | 87.04±0.54 | 87.34±0.37 | 87.3±0.38 | 74.38±0.85 | 93.87±0.36 |
| **Combined** | **89.61±0.16** | **89.84±0.29** | **89.38±0.14** | **89.43±0.13** | **79.23±0.33** | **95.52±0.12** |

**Figure 4.**
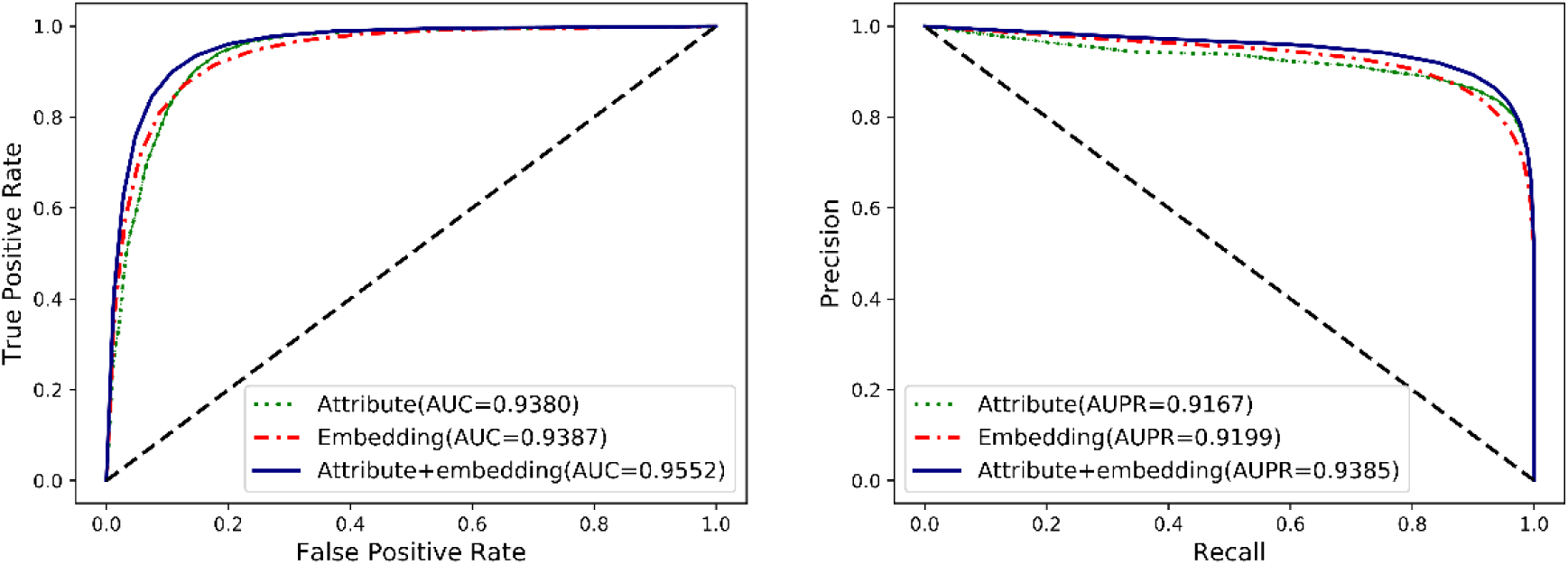
Comparison of nodes network embedding and attribute features.

The attribute of each type of node are obtained by the most widely used feature extraction methods in its related research, such as k-mer frequency for lncRNA, miRNA and protein sequences, fingerprint for drug chemical structure, and semantic similarity for disease phenotype. The network embedding of nodes are learned by the SDNE algorithm on the training set. In order to evaluate and compare the impact of each kind of feature on the final classification performance of the MAN-SDNE model, we performed our model separately using individual network embedding feature, attribute feature and combined use of these two features. As the results listed in Table 4 and Figure 4, the individual effects of network embedding features and attribute features are already acceptable, and the large-scale network representation features are at least as good as state-of-the-art attribute features. Moreover, the combination of these two features can achieve the best results, demonstrating the complementary potential of the two features.

### Comparison of widely used machine learning classifiers

In order to highlight the importance of constructing MAN under a systematic view, the most commonly used Random Forest model are choose as the classifier in the link prediction scenario. In this section, we will compare Random Forest with other widely used machine learning model, including Naïve Bayes (NB), Logistic Regression (LR), AdaBoost, and XGBoost, under the same experimental conditions. Both the Random Forest classifier and other contrast models use default parameters to avoid bias. The results are shown in Table 5 and Figure 5 as below.

**Table 5.** Compare with widely used machine learn classifiers.

| Method | Acc. (%) | Sen. (%) | Spec. (%) | Prec. (%) | MCC (%) | AUC (%) |
| --- | --- | --- | --- | --- | --- | --- |
| NB | 68.2±0.85 | 47.31±1.34 | 89.1±0.68 | 81.27±1.18 | 40.08±1.73 | 80.6±0.84 |
| LR | 80.65±0.98 | 82.02±1.02 | 79.28±2.65 | 79.88±1.88 | 61.35±1.9 | 88.03±1.25 |
| AdaBoost | 83.55±0.22 | 85.39±0.47 | 81.72±0.46 | 82.37±0.33 | 67.15±0.45 | 91.19±0.19 |
| XGBoost | 84.22±0.16 | 88.65±0.26 | 79.79±0.46 | 81.43±0.32 | 68.71±0.29 | 91.83±0.23 |
| **Proposed method** | **89.61±0.16** | **89.84±0.29** | **89.38±0.14** | **89.43±0.13** | **79.23±0.33** | **95.52±0.12** |

**Figure 5.**
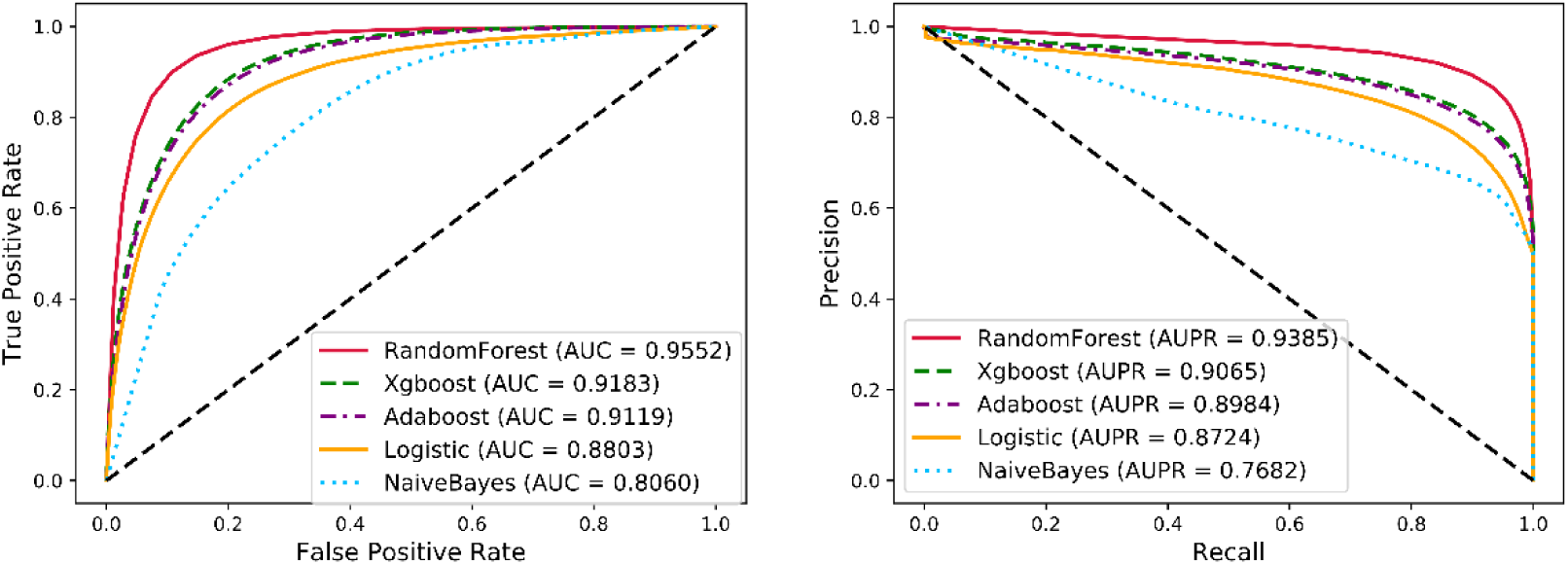
Comparison of widely used machine learning classifiers and MAN-SDNE.

The proposed method achieves the best results on all indicators with a high AUC of 95.52 and a high AUPR of 0.9385, while also having the smallest standard deviation. And the other machine learning classifiers also have good performance. NB can obtain better results when the properties of the samples are independent of each other. In this experiment, there are cases where the attributes are not independent and the cross-joins together affect the final classification effect. LR is essentially a linear classifier whose performance is limited by the distribution of data and does not perform well in this case. All classifier parameters are default, which may cause AdaBoost and XGBoost to be inappropriate or over-fitting on this task. The results verify the robustness of the Random Forest and the high discriminability of features.

### Case study: predicting lncRNA-protein interactions using MAN-SDNE

The effectiveness and robustness of the feature representation and classifier have been proved in the above experiments. In fact, MAN-SDNE can not only be used to predict the interaction between nodes under homogeneous conditions, but also play a role in the specific interaction types prediction in the network. In this section, we use MAN-SDNE to predict lncRNA-protein interactions. More specifically, in MAN dataset, there is 690 lncRNA-protein interaction, and the count of other associations is 104856, we divided the lncRNA-protein pairs into 5 equal subsets, according the strategy of five-fold cross-validation, prepare the train set and test set. In each fold of cross-validation, the training set is added to the remaining 104856 edges to train the SDNE for learning the network embedding of nodes. The conjoint triad (3-mer frequency) of lncRNA and protein sequence is used as node attribute. The results are shown in Figure 6. As shown in Figure 6A, MAN-SDNE can achieve good performance with an AUC of 80.28%. Meanwhile, we also compared the effects of attribute, network embedding for predicting lncRNA-protein interactions, their performances are shown in Figure 6B, Figure 6C, respectively. And the comparison of them is shown in Figure 6D. The results demonstrate the capability of MAN-SDNE to predict heterogeneous interaction types. It also proves that the cell is a complete unit of life, and the interaction of biomolecules in the cell is interconnected to maintain the normal conduct of life activities.

**Figure 6.**
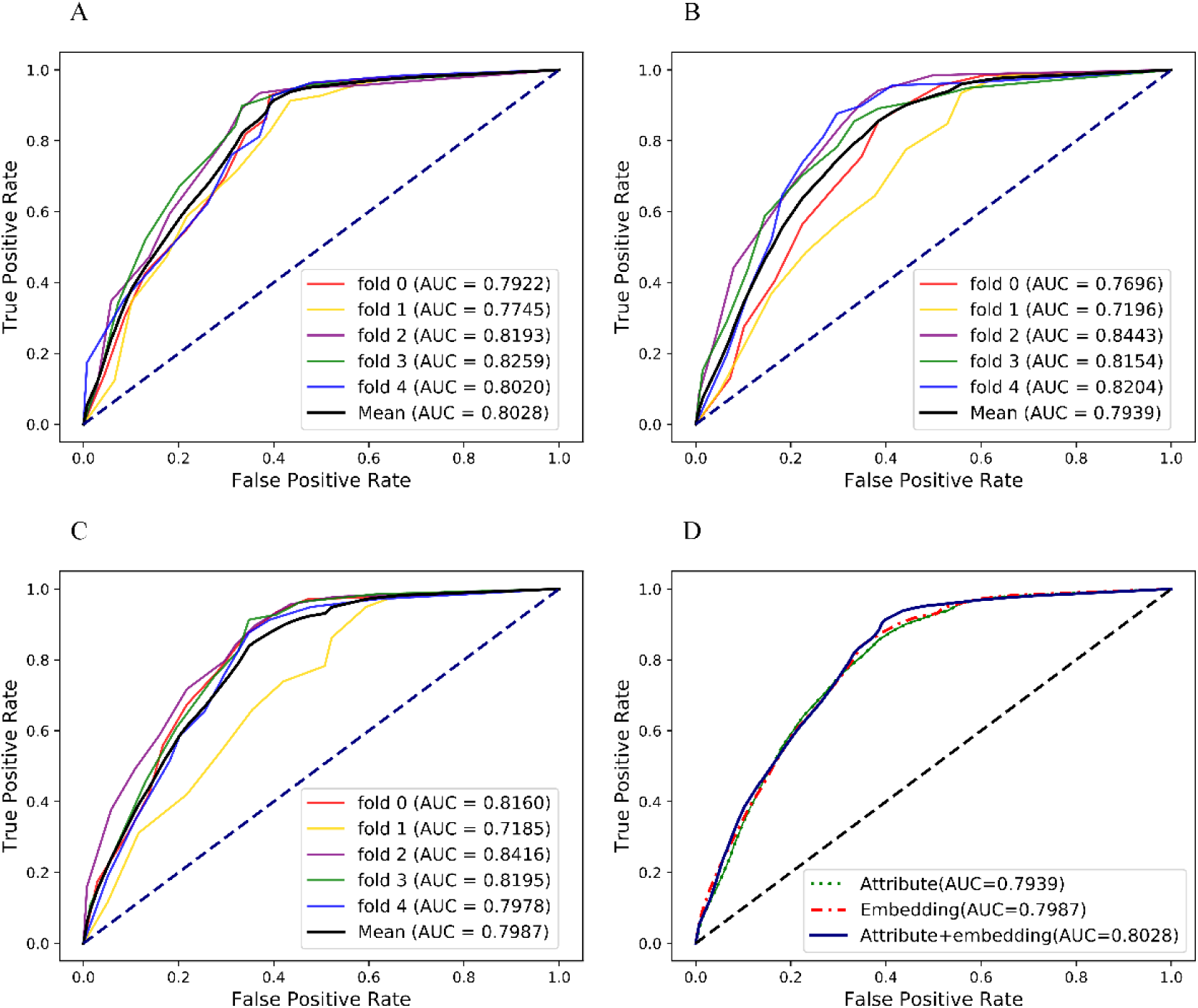
The ROC performance of MAN-SDNE for predicting lncRNA-protein interactions. (A) five-fold cross-validation performance using network embedding and attribute feature of nodes; (B) five-fold cross-validation performance using only nodes attribute feature; (C) five-fold cross-validation performance using only network embedding feature; (D) performance comparison of different features.

## Conclusion

In this study, we revealed and defined a molecular associations network of various intermolecular associations. The difference from previous studies is that we consider these intermolecular interactions as interconnected entities to understand the wide-ranging interactions between molecules in human cells from a systematic view. The SDNE algorithm is applied to learn network embedding of nodes from the MAN. The k-mer frequency, fingerprint, and semantic similarity are also employed as the attribute of nodes. The Random Forest classifier is trained for link prediction. And we analyzed and compared the features, models and results. In addition, we did a case study using MAN-SDNE to predict lncRNA-protein interactions, which indicate the ability of MAN-SDNE to predict specific types of interactions in entire molecular associations network. Above these, for diseases, microbes, objects that are difficult to directly represents, MAN network embedding can be regarded as a good method of providing discriminative feature representation. In this work, different types of edge are treated as homogeneous, the node attributes are directly combined to the embedding, and some known types of interactions have not yet merged into our network, which can be improved in our future work.

## Author Contributions

H-C. Y conceived the algorithm, carried out analyses, prepared the data sets, carried out experiments, and wrote the manuscript; Z-H. Y and Z-H. G designed, performed and analyzed experiments and wrote the manuscript; All authors read and approved the final manuscript.

## Acknowledgments

This work is supported by the National Natural Science Foundation of China, under Grants 61572506, in part by the NSFC Excellent Young Scholars Program, under Grants 61722212, in part by the Pioneer Hundred Talents Program of Chinese Academy of Sciences.

## Conflicts of interest

The authors declare no conflict of interest.

